# The Dlg-module and clathrin-mediated endocytosis regulate EGFR signaling and cyst cell-germline coordination in the *Drosophila* testis

**DOI:** 10.1101/419937

**Authors:** Fani Papagiannouli, Cameron Wynn Berry, Margaret T. Fuller

**Affiliations:** Department of Developmental Biology, Beckman Center, Stanford University School of Medicine, CA 94305-5329, Stanford, USA; Institute for Genetics, University of Cologne, Cologne, 50674, Germany

**Keywords:** *Drosophila* testis, germline, soma-germline communication, spermatogenesis, Dlg, Scrib, Lgl, Shibire, AP-2α, endocytosis, polarity, EGFR, signaling regulation, clathrin-mediated endocytosis

## Abstract

Tissue homeostasis and repair relies on proper communication of stem cells and their differentiating daughters with the local tissue microenvironment. In the *Drosophila* male germline adult stem cell lineage, germ cells proliferate and progressively differentiate enclosed in supportive somatic cyst cells, forming a small organoid, the functional unit of differentiation. Here we show that cell polarity and vesicle trafficking influence signal transduction in cyst cells, with profound effects on the germ cells they enclose. Our data suggest that both the cortical components Dlg, Scrib, Lgl and the clathrin-mediated endocytic (CME) machinery downregulate EGFR signaling. Knockdown of *dlg, scrib, lgl* or CME components in cyst cells resulted in germ cell death, similar to increased signal transduction via the EGFR, while lowering EGFR or downstream signaling components rescued the defects. This work provides new insights on how cell polarity and endocytosis cooperate to regulate signal transduction and sculpt developing tissues.

## INTRODUCTION

The genesis and maintenance of functional tissues and organs require close communication between disparate cell types, with exchange of short-range signals regulating cell proliferation and survival, cell fate, and local patterning. Tissues or highly differentiated cell types with a high turnover rate such as intestinal epithelium, red blood cells, or skin rely on adult stem cells for constant replenishment of differentiated cell populations (Monahan and Starz-Gaiano, 2016). Similarly, habitually quiescent adult stem cells are maintained in reserve for repair of tissue damage or response to physiological conditions such as puberty, pregnancy, or nutritional state (Merrell and Stanger, 2016). As during organogenesis, tissue homeostasis and repair by adult stem cell lineages rely on proper short-range communication with the local microenvironment, and equilibrium of stem cell maintenance vs. differentiation (Losick et al., 2011; Matunis et al., 2012; Papagiannouli, 2014; Papagiannouli and Lohmann, 2012).

In *Drosophila*, male germline stem cells (GSCs) reside at the apical tip of the testis, flanked by somatic cyst stem cells (CySCs). Both the germline and the somatic stem cells are maintained through their association with the hub, a cluster of normally non-dividing somatic cells forming the niche organizer (Fig.1A). By asymmetric cell division with an oriented spindle, each GSC produces a new GSC attached to the hub plus a distally located gonialblast (GB). The gonialblast initiates four rounds of transit-amplifying (TA) mitotic divisions with incomplete cytokinesis, giving rise to 2-, 4-, 8-and eventually 16-interconnected germ cells. Immediately after reaching the 16-cell stage, the germ cells undergo a final round of DNA synthesis, enter meiotic prophase, and turn on the spermatocyte transcription program for meiosis and spermatid differentiation. The somatic CySCs also divide asymmetrically, producing CySCs that remain associated with the hub plus distally located post-mitotic daughters that become cyst cells (CC) (Fuller and Spradling, 2007). Two somatic cyst cells enclose each gonialblast, forming a two-cell squamous epithelium that encases the mitotic and meiotic progeny of that gonialblast throughout the rest of male germ cell differentiation until sperm individualization. Notably, the cyst cells codifferentiate with the germ cells they enclose (Gonczy and DiNardo, 1996), and are required for germ cell differentiation (Lim and Fuller, 2012).

**Figure 1:**
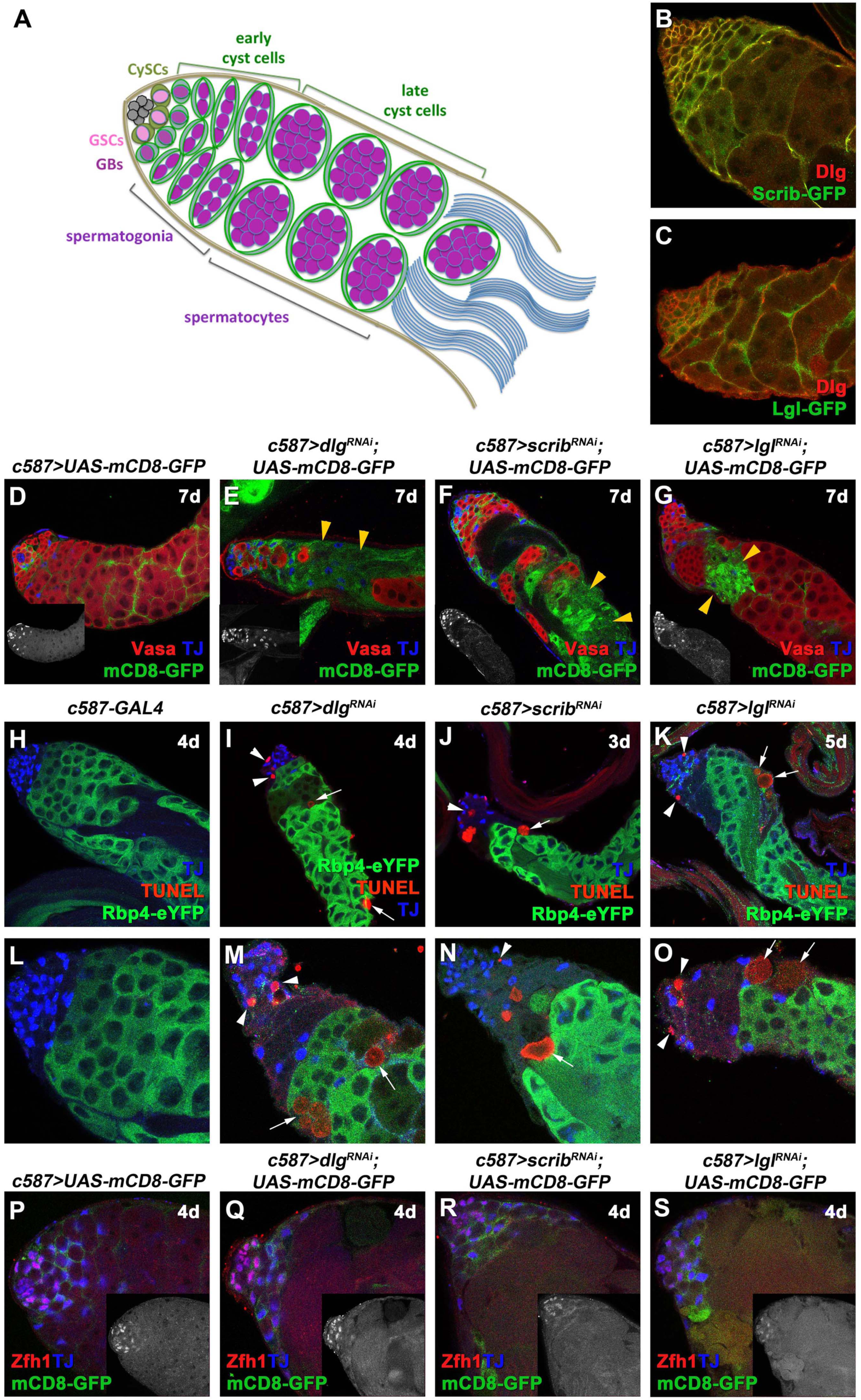
Knockdown of *dlg, scrib* or *lgl* function in cyst cells leads to cell non-autonomous germ cell death. **(A)** Diagram of early spermatogenesis in *Drosophila*. GSC: germline stem cell, GB: gonialblast, CySC: somatic cyst stem cell. **(B, C)** Adult testes from flies with endogenously tagged (B) Scrib-GFP and (C) Lgl-GFP immunostained stained for Dlg (red) and GFP (green). Adult testes of the indicated genotypes in the background of *Gal80*^*ts*^. **(D-G)** *mCD8-GFP* (green) expressed in cyst cells, anti-Vasa (red; germline), anti-TJ (blue; early cyst cells). Yellow arrowheads: mCD8-positive cyst cell regions. **(H-O)** anti-TJ (blue), GFP (green; Rbp4-GFP marking spermatocytes) and TUNEL (red) marking apoptotic double-strand breaks. (L-O) higher magnifications of testes apical regions from (H-K). White arrowheads: dying spermatogonia, White arrows: dying spermatocytes. **(P-S)** *mCD8-GFP* (green) expressed in cyst cells, anti-Zfh1 (red; CySC and immediate daughters) and anti-TJ (blue). Small inset pictures in (P-S) show the anti-Zfh1 staining only. Newly eclosed male flies were shifted at 30°C to activate the RNAi for 3, 4, 5 or 7 days (d). Testes oriented with anterior at left. Image frame: (D-K) 246μ m, (B, C, L-S) 123μ m. This figure is associated with **Figures S1, S2, S3**.

In addition to physically enclosing each packet of mitotic sister germ cells throughout spermatogenesis, the somatic cyst cells control spermatogenesis from initial differentiation to mature sperm production (Leatherman, 2013; Zoller and Schulz, 2012). Signals from the germline to the cyst cells via the Epidermal Growth Factor play a major role (Hudson et al., 2013; Kiger et al., 2000; Sarkar et al., 2007; Schulz et al., 2002; Tran et al., 2000). Activation of the EGFR and its downstream signal transduction pathway, leading to phosphorylation of MAPK in the somatic cyst cells, is required for germ cells to properly enter and execute the mitotic proliferation program of synchronous transit-amplifying divisions that is the first step of differentiation (Kiger et al., 2000; Sarkar et al., 2007; Schulz et al., 2002; Tran et al., 2000). Expression of the EGF class ligand *vein* is subsequently induced in cyst cells (Singh et al., 2016) to consolidate and enforce EGFR signaling cell-autonomously as has been shown in other tissues (Yarnitzky et al., 1998). Activation of the EGFR on cyst cells at even higher levels is required for germ cells to exit the mitotic proliferation program and switch to the spermatocyte state (Hudson et al., 2013).

Here we show that function of the cell polarity components *discs large (dlg), scribble (scrib)* and *lethal (2) giant larvae (lgl)* in the cyst cell lineage are crucial for proper formation of a functional cyst microenvironment that supports the survival of differentiating germ cells. *dlg, scrib* and *lgl* were first identified as tumor suppressor genes whose loss of function leads to neoplastic transformation (Bilder and Perrimon, 2000; Donohoe et al., 2018; Elsum et al., 2012; Goode and Perrimon, 1997; Li et al., 2001; Mechler et al., 1985; Woods et al., 1996). Dlg, Scrib and Lgl, collectively called the Dlg-module, are highly conserved polarity and scaffolding proteins involved in: (i) establishment and maintenance of apical/basal cell polarity in columnar epithelia in cooperation with the Par-(Bazooka/Par3, Par6, αPKC) and Crumbs-polarity complexes, (ii) vesicle and membrane trafficking in *Drosophila*, yeast and mammals, and (iii) cooperation with signaling pathways (e.g. Ras, Notch and JNK signaling pathways) in normal tissues and in cancer (de Vreede et al., 2014; Elsum et al., 2012; Gui et al., 2016; Parsons et al., 2014; Roegiers et al., 2005; Uhlirova and Bohmann, 2006). Previous work has shown that somatic cells fail to extend projections and encapsulate the germ cells in embryonic gonads of male flies mutant for *scrib* or *dlg*, suggesting a role in establishing intimate germline-soma contacts (Marhold et al., 2003; Papagiannouli, 2013). Larval and adult males mutant for *dlg, scrib*, or *lgl* had extremely small testes with reduced number of GSCs, accumulation of cyst cells, impaired germ cell differentiation, resulting in sterility (Fairchild et al., 2017; Papagiannouli, 2013; Papagiannouli and Mechler, 2009). Importantly, expression of a *dlg* transgene in cyst cells of *dlg* mutant larval testes rescued encapsulation of the germline by somatic cells and restored the CySC-GSC arrangement around the niche, and the architecture and integrity of spermatogonial and spermatocyte cysts (Papagiannouli and Mechler, 2009).

Our results suggest that the highly conserved Dlg-module cooperates with clathrin-mediated endocytic (CME) components to downregulate the EGFR signaling in somatic cyst cells. We show that the cell-type specific knockdown of the Dlg-module or CME components in cyst cells results in increased levels of downstream mediators of EGFR signaling in cyst cells, accompanied by non-autonomous germ cell death, phenocopying the effect of EGFR overactivation in cyst cells. Lowering the levels of EGFR signal transduction components in somatic cyst cells partially rescued the observed defects and restored germ cell survival in animals in which function of Dlg-module or clathrin-mediated endocytic components had been knocked down in cyst cells. Consequently, we conclude that the Dlg-module and clathrin-mediated endocytosis normally down-regulate signaling via the EGFR and that this fine-tuning is critical for germline cysts to differentiate gradually and proceed through subsequent developmental steps.

## RESULTS

### Knockdown of *dlg, scrib* or *lgl* function in cyst cells leads to germ cell death

Dlg, Scrib and Lgl are coexpressed in early cyst cells that encapsulate gonialblasts and spermatogonia, and in late cyst cells that enclose spermatocyte cysts (Fig.1B, 1C). Function of Dlg-module components was impaired in the cyst cell lineage using the *c587-GAL4* driver with *UAS-gene*^*RNAi*^ lines to knockdown expression of *dlg, scrib* or *lgl*. The flies also carried an *αtubGal80*^*ts*^ transgene, which blocks activity *of GAL4* at 18°C but not at 30°C, allowing acute downregulation of gene function in adults, after normal testis anatomy had been set up. Analysis of the tissue 2, 4 or 7 days after the shift to 30°C allowed us to observe progression of the phenotypes over time (e.g. Fig.S1) and avoiding confusion with effects on testis architecture due to earlier roles in development.

Knockdown of *dlg, scrib* or *lgl* function in somatic cyst cells had non autonomous effects on differentiating germ cells, resulting in loss of late spermatogonia or early spermatocytes. The germ cell loss was visible as a gap in Vasa staining (Fig. 1E-1G) that became progressively larger with longer exposure to knockdown conditions (Fig. S1A-S1L), while in control testes Vasa marked all densely packed germ cells (Fig. 1D). As this germ cell loss was likely caused by apoptosis, we used the TUNEL assay to label double-strand DNA breaks in apoptotic cells. TUNEL staining revealed spermatocytes, marked by expression of an Rbp4-GFP transgene, undergoing apoptosis in testes in which *dlg, scrib* or *lgl* had been knocked down in the cyst cell lineage (Fig. 1H-1O).

Even after 7 days at 30°C, knockdown testes typically retained early spermatogonia that, based on staining for phosphorylated Histone 3 (PH3), retained their capacity to proliferate (Fig. S2A-S2H) and produced at least some cysts of germ cells positive for the late spermatogonial marker Bam (Fig. S2I-S2P’). After several days of *dlg, scrib* or *lgl* knockdown in cyst cells (when spermatocytes formed prior to the knockdown had presumably progressed to differentiation) only few Rbp4-GFP marked spermatocyte cysts remained (Fig. S3H-S3J).

The loss of germ cells was accompanied by clustering of the *dlg, scrib* or *lgl* depleted cyst cells in the affected regions, manifested as patches of the membrane CD8-GFP (mCD8-GFP) marker expressed in cyst cells (Fig.1D-1G; yellow arrowheads). Knockdowns performed for 2 days (2d) revealed that the defects appeared initially around TJ-positive early cyst cells that clustered together as germ cells first began to be eliminated, near the spermatogonia/spermatocyte boundary (Fig. S1A-S1D). The mCD8-positive patches progressively expanded (Fig. S1E-S1L; yellow arrowheads) as germ cell loss continued in flies incubated at 30°C for 4 and 7 days. Double knockdown of *dlg* with *scrib* or *lgl* led to phenotypes similar to the single knockdowns (Fig.S1M-S1P), in agreement with the Dlg-module proteins acting in the same pathway.

Consistent with the continued presence of early spermatogonia at the testis apical tip, knockdown of *dlg, scrib* or *lgl* function in the cyst cell lineage did not appear to drastically alter cyst cell identity. Immunostaining for Zfh1, which marks CySCs and their immediate daughter cyst cells (Leatherman and Dinardo, 2008), showed Zfh1 positive nuclei clustered at the apical tip of testes, as in wild type (Fig.1P-1S). Likewise, cyst cells that had lost the Zfh1 marker but still expressed the early cyst cell marker TJ were ranked a bit farther from the hub than the Zfh1+ cyst cells in the knockdown testes, as in wild type (Fig. 1P1-1S).

### The effects of loss of Dlg, Scrib or Lgl resemble the phenotypes of the EGFR pathway overactivation

The phenotype resulting from suppression of *dlg, scrib* or *lgl* function in cysts cells was strongly reminiscent of the overactivation effects of constitutively active EGFR (EGFR^CA^) [(Hudson et al., 2013) and Fig. 2C] or *Ras85D* (*UAS-Ras*^*V*^*12*^^) (Fig. 2D) in cyst cells. Immunostaining of adult testes in which either *UAS-EGFR*^*CA*^ or *UAS-Ras85D*^*V*^*12*^^ was overexpressed in cyst cells for 7 days and 2 days respectively, revealed that early germ cells near the testis tip persisted, while some Bam-positive late spermatogonial cysts were still formed (Fig.S3E-S3G’), which retained their proliferation capacity (Fig. S2J, S2J’). Slightly farther from the testis tip, cyst cells lacking associated germ cells accumulated, visible as clusters of TJ positive nuclei embedded in patches of mCD8-GFP (Fig. 2C, 2D; yellow arrowheads). TUNEL staining of testes 3 days after forced expression of *UAS-EGFR*^*CA*^ showed dying germ cells within the Rbp4-positive spermatocyte region (Fig. 2G), similar to that observed after knockdown of Dlg-module components in cyst cells (Fig. 2F; Fig. 1I-1K). Upon longer incubation under knockdown conditions, few Rbp4-positive spermatocytes remained (Fig.S3K, S3K’). Importantly, overexpression of a wild type form of EGFR or of *Ras85D* in cyst cells resulted in milder phenotypes (Fig. S4A-S4C) reminiscent of the defects observed upon short-term (2 days) inhibition of *dlg, scrib* or *lgl* (compare Fig.S1A-S1L with Fig.S4A-S4C). Strikingly, knockdown of *dlg, scrib, lgl* in cyst cells resulted in elevated levels of phosphorylated MAPK (dpERK), a canonical downstream effector of the EGFR signal transduction pathway. In testes co-stained for dpERK and the early cyst cell marker TJ, cyst cells depleted for *dlg* (Fig. 2I), *scrib*, or *lgl* (Fig. S4D, S4F) showed increased dpERK levels in TJ-positive cyst cell nuclei compared to nuclei of similarly staged cyst cells in control testes (Fig. 2H, 2K). Similar increase in nuclear dpERK was observed in cyst cells in which *EGFR*^*CA*^ was overexpressed for 4 days (Fig. 2J, 2K).

**Figure 2:**
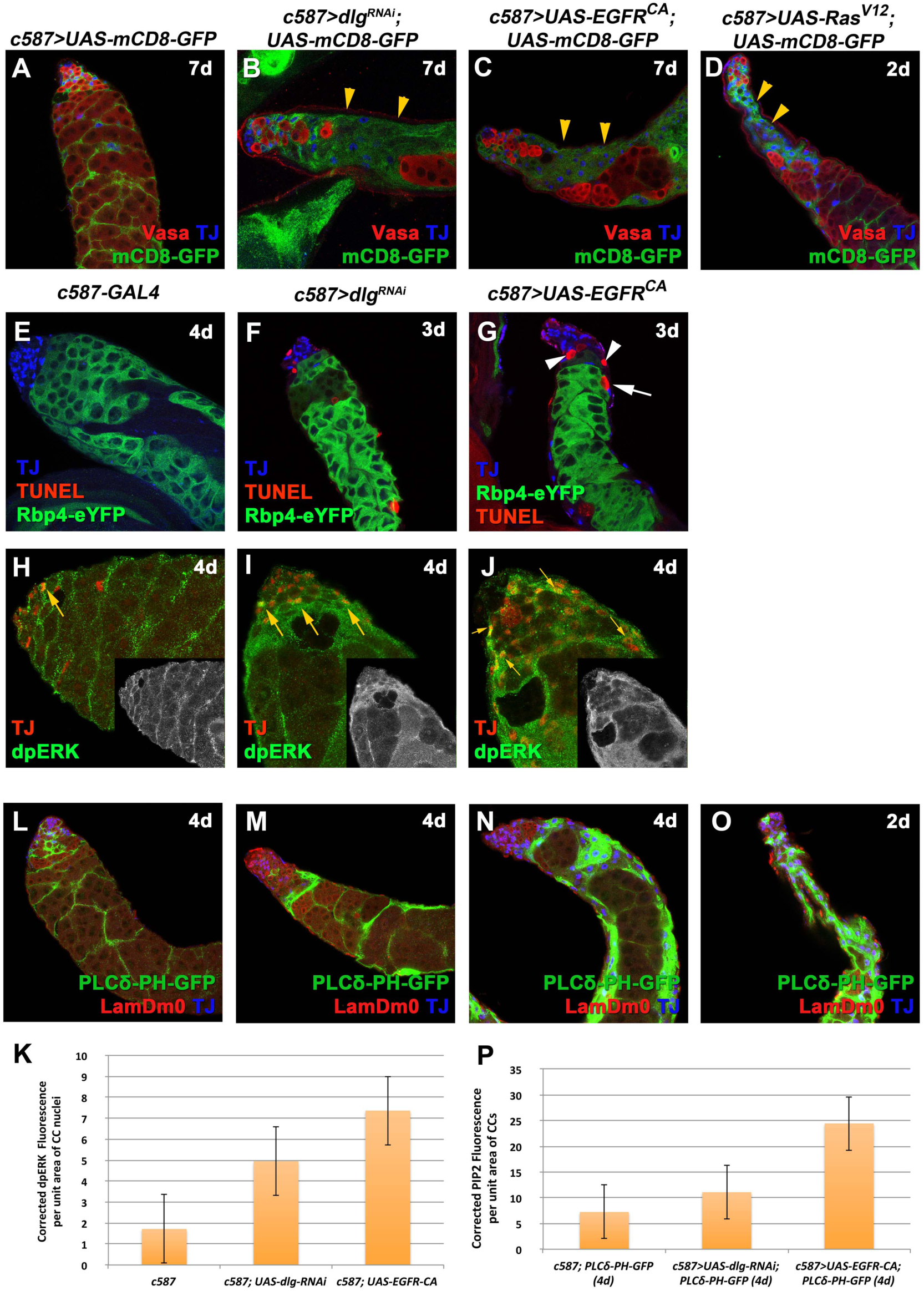
Knockdown of *dlg, scrib* or *lgl* and overactivation of EGFR receptor signaling in cyst cells have similar phenotypes. Adult testes of the indicated genotypes in the background of the *Gal80*^*ts*^. **(A-D)** *mCD8-GFP* (green) expressed in cyst cells; anti-Vasa (red; germline) and anti-TJ (blue; early cyst cells). Yellow arrowheads: mCD8-positive cyst cell regions. **(E-G)** anti-TJ (blue), GFP (green; Rbp4-GFP marking spermatocytes) and TUNEL (red) marking apoptotic double-strand breaks. White arrowheads: dying spermatogonia. White arrows: dying spermatocytes. **(H-J)** anti-TJ (red) and anti-dpERK (green). Yellow arrows: examples of cyst cells double-labelled for TJ and dpERK. Small inset pictures show the anti-dpERK staining only. **(K)** Quantification of corrected fluorescent dpERK levels in cyst cell (CC) nuclei depleted of *dlg* function or overexpressing a constitutively active (CA) EGFR. **(L-O)** the PIP2 reporter PLCd-PH-GFP expressed in cyst cells (green), anti-LamDm0 (red) marking cyst cells and early germ cells (GSCs, GBs and spermatogonia) and anti-TJ (blue; early cyst cells). **(P)** Quantification of corrected fluorescent PIP2 levels in cyst cells (CC) depleted of *dlg* function or overexpressing a constitutively active (CA) EGFR. Newly eclosed male flies were shifted at 30°C to activate the RNAi for 2, 3, 4 or 7 days. Testes oriented with anterior at left. Image frame: (A-G, H-J) 246μ m, (L-O) 123μ m This figure is associated with **Figure S4**

As EGFR activation can also result in elevated levels of the membrane phospholipid Phosphatidylinositol 4,5-bisphosphate [PtdIns(4,5)P2] (PIP2) (Czech, 2000), we assessed PIP2 levels in cyst cells by expressing the PLCδ-PH-GFP reporter, which contains the PIP2-specific pleckstrin-homology domain of phospholipase Cδ (Gervais et al., 2008). Levels of the PIP2 reporter (Fig. 2P, S4K), appeared higher in testes in which cyst cells were depleted for *dlg* (Fig. 2M), *scrib* or *lgl* function (Fig. S4H, S4I) compared to control testes (Fig.2L). Similar higher levels of the PIP2 reporter were also observed in testes in which *EGFR*^*CA*^ or *Ras*^*V*^*12*^^ were overexpressed in the cyst cell lineage (Fig. 2N, 2O).

### Lowering levels of EGFR signal transduction pathway components in cyst cells partially rescues *dlg, scrib* and *lgl* knockdown phenotypes

Given the enhanced dpERK activity following *dlg, scrib* or *lgl* inhibition, we tested if reduced EGFR signaling might rescue the *dlg*-module loss of function phenotypes. Strong loss of EGFR function results in failure of cyst cells to encapsulate early spermatogonia and defects in the ability of germ cells to properly enter the transit-amplifying program of four rounds of synchronous mitotic divisions (Kiger et al., 2000; Sarkar et al., 2007; Schulz et al., 2002; Tran et al., 2000). As a result, early spermatogonia of mixed identity proliferate extensively and the cyst cells are pushed to the testis periphery. To circumvent the extreme early phenotype caused by a complete EGFR loss, we took three different approaches to interfere with the EGFR signaling. We lowered the gene dose by using flies heterozygous for the null *EGFR*^*K*^*05115*^^ allele (*EGFR*^*K*^*05115*^^/+) (Fig.3C, S5D1-3), we overexpressed a dominant negative EGFR allele (*UAS-EGFR*^*DN*^) (Fig.4B), or finally used transgenic RNAi to knockdown the EGFR (*UAS-EGFR*^*RNAi*^) or Ras (*UAS-Ras*^*RNAi*^) in the background of *dlg, scrib* and *lgl* knockdowns in cyst cells.

**Figure 3:**
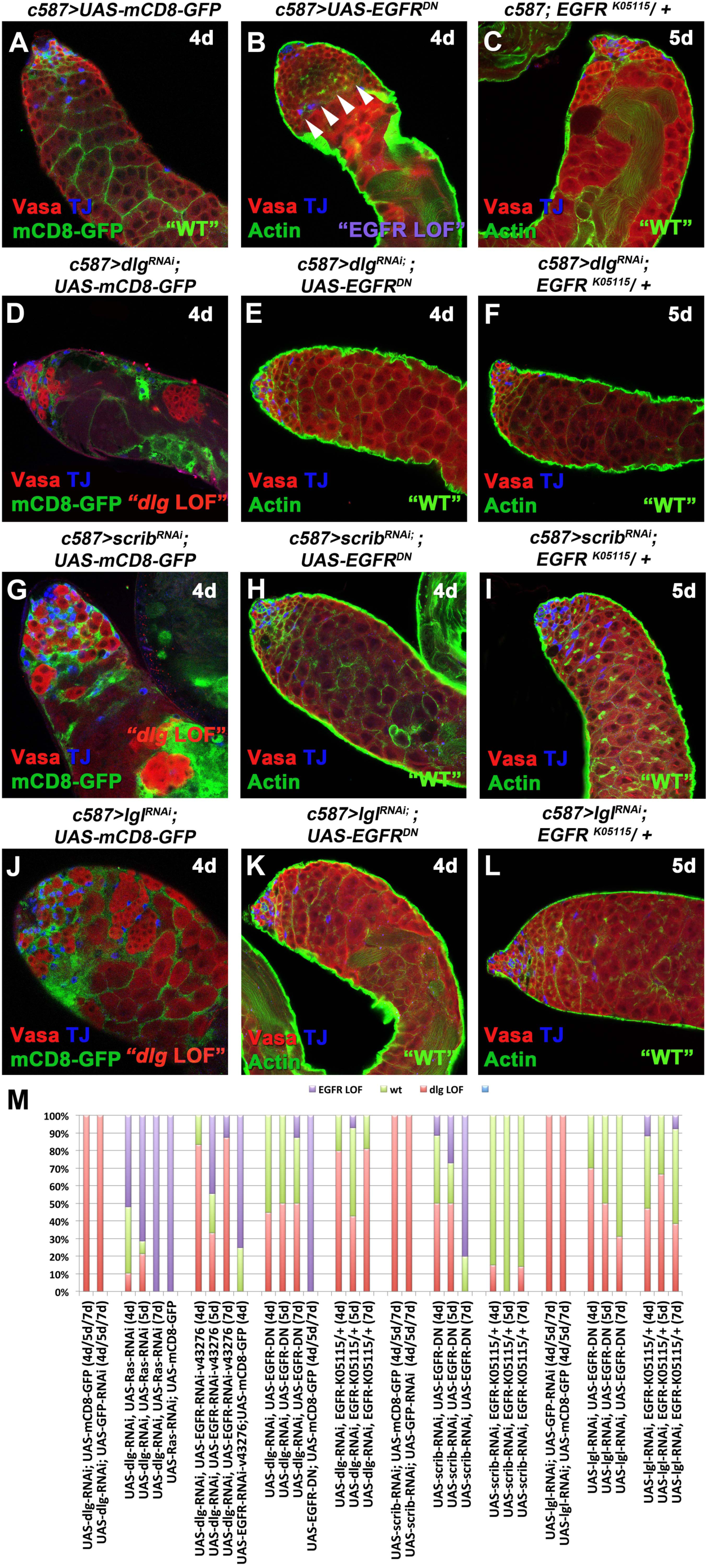
Lowering EGFR signaling levels can rescue the *dlg, scrib* and *lgl* loss of function phenotypes in cyst cells. Adult testes of the indicated phenotypes in the background of the *Gal80*^*ts*^. **(A, D, G, J)** *mCD8-GFP* (green) expressed in cyst cells; anti-Vasa (red; germline); anti-TJ (blue; early cyst cells), **(B, C, E, F, H, I, K, L)** Actin stained for phalloidin (green; cyst cells and germline fusome); anti-Vasa (red); anti-TJ (blue). (B) White arrows indicate overproliferating early spermatogonia upon EGFR loss of function. **(M)** Quantifications of the different phenotypic classes accompanying each genotype, organized in order of phenotypic strength: weak and strong “dlg LOF”, “wt” and “EGFR LOF”. Newly eclosed male flies were shifted at 30°C to activate the RNAi for 4, 5 and 7 days. Testes oriented with anterior at left. Image frame 246μ m. This figure is associated with **Figure S5**

**Figure 4:**
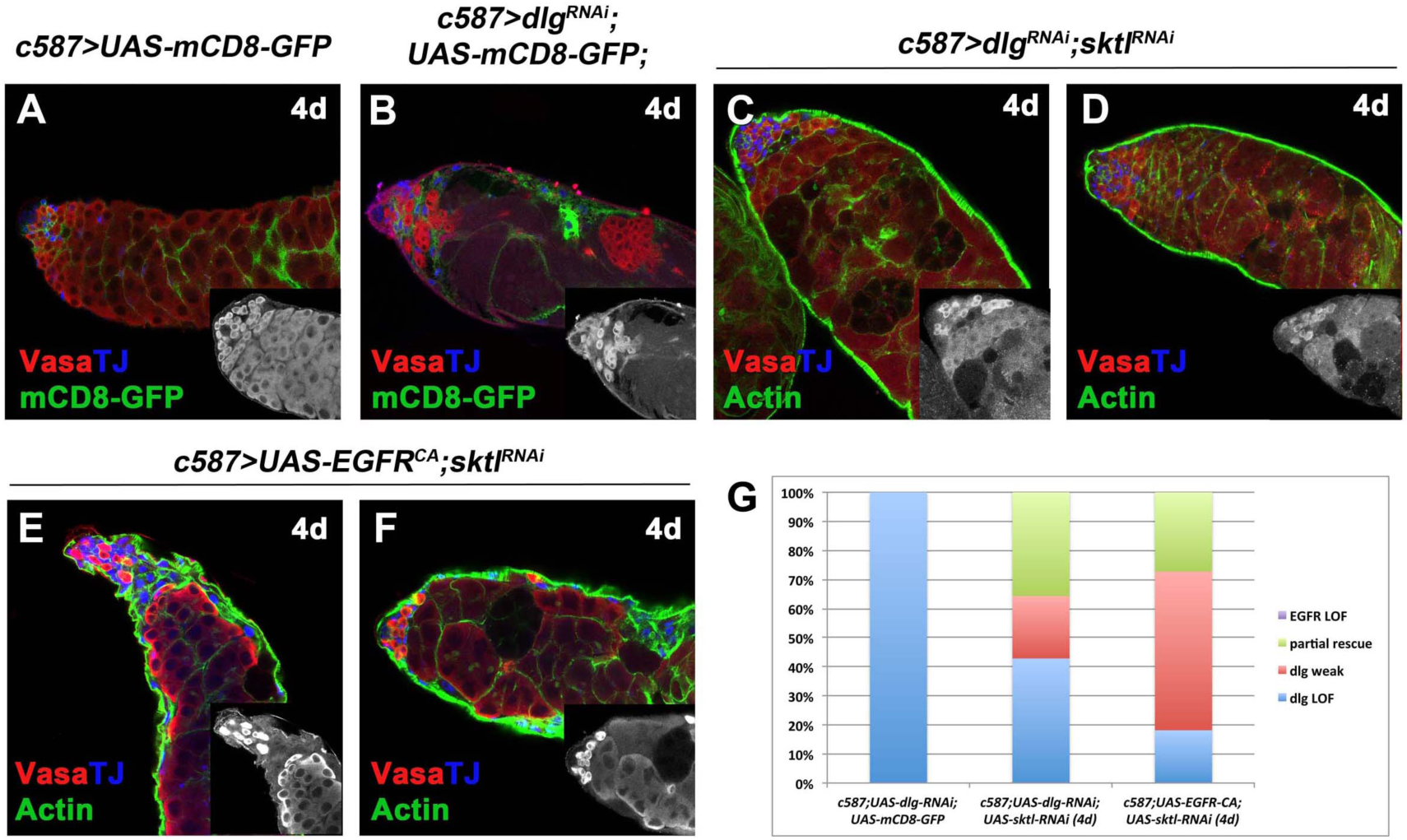
Knockdown of the dPIP5K kinase *sktl* in cyst cells can partially rescue the *dlg* loss of function or EGFR overactivation phenotypes. Adult testes of the indicated phenotypes in the background of the *Gal80*^*ts*^. **(A, B)** *mCD8-GFP* (green) expressed in cyst cells; anti-Vasa (red; germline); anti-TJ (blue; early cyst cells), **(C-F)** F-actin stained for phalloidin (green; cyst cells and germline fusome); anti-Vasa (red); anti-TJ (blue). **(G)** Quantifications of the different phenotypic classes accompanying each genotype, organized in order of phenotypic strength: “dlg LOF”, “dlg weak” and “partial rescue”. No testes with “EGFR LOF” phenotypes emerged. Newly eclosed male flies were shifted at 30°C to activate the RNAi for 4 days. Testes oriented with anterior at left. Image frame 246μ m.

To control for possible effects of multiple UAS constructs, limiting the effectiveness of the *GAL4* driver or multiple RNAi constructs limiting the effectiveness of the RNAi machinery, control flies were set up to carry the same number of *UAS* transgenes using two types of controls: *UAS-mCD8-GFP* or *UAS-eGFP-RNAi*. So for example, *c587>UAS-dlg*^*RNAi*^; *UAS-EGFR*^*DN*^ flies were compared to control flies of the genotype *c587>UAS-dlg*^*RNAi*^; *UAS-mCD8-GFP*, while the best control for *c587>UAS-dlg*^*RNAi*^; *UAS-EGFR-RNAi* would be *UAS-dlg*^*RNAi*^; *UAS-GFP*^*RNAi*^. All genetic combinations carried the *Gal80*^*ts*^ transgene. Because different *UAS* transgenes have different expression strengths, rescue experiments were performed by shifting flies to 30°C for 4, 5 and 7 days, followed by phenotypic classification.

The observed phenotypes were classified into the following categories: 1) “dlg LOF” included testes that mimicked the effect of acute *dlg* cyst cell knockdown with strong decrease in number of germ cells (comparable to Fig.1E, S1F, S1J) but also very weak knockdowns where only small patches of cyst cell clusters devoid of germ cells were identified (comparable to Fig. S1B-S1D); 2) “wt” for testes with restored spermatogonial and spermatocyte cysts similar to wild type; 3) “EGFR LOF” for all phenotypes that resulted in overproliferation of early spermatogonia resembling strong loss of EGFR signaling. Representative examples of the different phenotypic classes, reflecting the variability in strength and penetrance of “dlg LOF” and “EGFR LOF” phenotypic classes in comparison to “wt” for the different genotypes are shown in Fig.3 and Fig.S5.

Although a range of phenotypes resulted from our rescue strategy, the percent of testes showing a “dlg LOF” phenotype was substantially reduced, usually to less than 50%, by lowering EGFR function in cyst cells in which Dlg-module components had been knocked down (Fig. 3, Fig. S5). In many cases 50% or more of the testes scored showed packed spermatogonial and spermatocyte cysts, and restored architecture as in wild type. With longer times of exposure to knockdown conditions (7 days), the strong “EGFR LOF” phenotype of overproliferating spermatogonial cysts came to predominate in some genotypes (Fig. 3M). Lowering the function of MAPK, encoded by the *Drosophila rolled (rl)* gene, also partially rescued *dlg* module loss of function phenotypes in cyst cells (Fig.S6).

Cyst cell knockdown of the *Drosophila skittles (sktl)* gene, encoding the dPIP5K kinase that synthesizes PIP2 from PI4K (Balakrishnan et al., 2015; Gervais et al., 2008), reduced the severity of both the *dlg* knockdown and *EGFR*^*CA*^overexpression phenotypes (Fig. 4). The dramatic germline apoptosis observed in cyst cells upon *dlg* knockdown (Fig. 4C, 4D) or EGFR overexpression (Fig. 4E, 4F) was reversed in many of the testes scored, and staining for filamentous actin (F-actin) revealed that cyst cells could encapsulate again groups of germ cells containing branched fusomes. However, knockdown of *sktl* could not completely rescue the phenotypes observed, as late spermatogonial cysts were not restored (Fig.4C-4F; inset pictures).

### Loss of Clathrin-mediated endocytosis in cyst cells causes defects resembling loss of Dlg-complex components

The amplitude and specificity of signaling levels is generally regulated by endocytosis of activated receptors that can either recycle back to the cell surface or get transported to lysosomes for degradation (Dobrowolski and De Robertis, 2011). EGFR becomes endocytosed mainly by clathrin-mediated endocytosis (CME) (Conte and Sigismund, 2016).

Given the significance of CME in EGFR signaling attenuation, we investigated how loss of CME components in cyst cells will affect EGFR signaling levels. Knockdown of CME components *shibire/shi* (the *Drosophila* homologue of *dynamin*), *AP-2α* (also known as *α-Adaptin)* or *Clathrin heavy chain (Chc)* in cyst cells, recapitulated the *dlg, scrib*, or *lgl* cyst cell knockdown and EGFR pathway overactivation phenotypes (Fig.5). As seen after knockdown of Dlg-module components, testes in which *shi, AP-2α* or *Chc* was depleted in the cyst cell lineage retained Vasa positive germ cells at the testis tip (Figure 5B-D), produced some Bam-positive spermatogonial cysts containing proliferating cells (Fig.S8), and had Zfh1 positive and TJ positive/Zfh1 negative cyst cell nuclei ranked at the testis apical tip, suggesting that the identity of early cyst cells remained intact and comparable to wild type controls (Fig. S7K-S7N’). However, away from the testis tip, testes depleted for *shi, AP-2α* or *Chc* function showed cyst cell nuclei positive for TJ clustered in mCD8-GFP positive regions devoid of the germ cells they normally encapsulate (Fig. 5A-5D), and germ cell death at the late spermatogonia/early spermatocyte boundary. TUNEL staining confirmed that spermatocytes expressing Rbp4-GFP underwent cell death (Fig.5E-5L).

**Figure 5:**
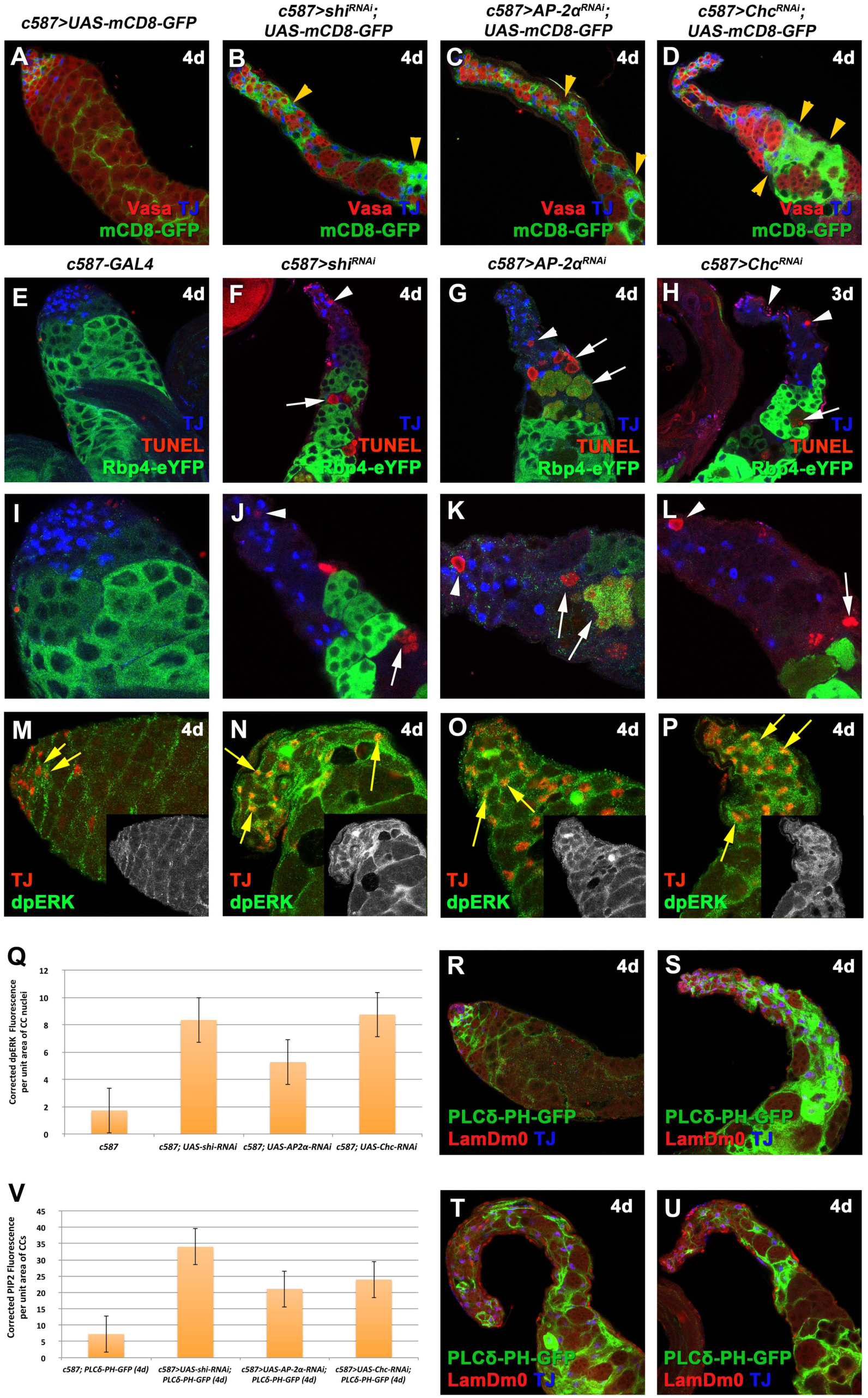
Defects in clathrin-mediated endocytosis show similar phenotypes as loss of function of Dlg-module components or EGFR overactivation in cyst cells. Adult testes of the indicated phenotypes in the background of the *Gal80*^*ts*^. **(A-D)** *mCD8-GFP* (green) expressed in cyst cells; anti-Vasa (red; germline) and anti-TJ (blue; early cyst cells), Yellow arrowheads: mCD8-positive cyst cell regions. **(E-L)** anti-TJ (blue), GFP (green; Rbp4-GFP marking spermatocytes) and TUNEL (red) marking apoptotic double-strand breaks. White arrowheads: dying spermatogonia. White arrows: dying spermatocytes. (I-L) higher magnifications of testes apical regions from (E-H). **(M-P)** anti-TJ (red) and anti-dpERK (green). Yellow arrows: examples of cyst cells double-labelled for TJ and dpERK. Small inset pictures show the anti-dpERK staining only. **(Q)** Quantification of corrected fluorescent dpERK levels in cyst cell (CC) nuclei depleted of *shi, AP-2α* or *Chc* function. **(R-U)** the PIP2 reporter PLCd-PH-GFP expressed in cyst cells (green), anti-LamDm0 (red) marking cyst cells and early germ cells (GSCs, GBs and spermatogonia), anti-TJ (blue; early cyst cells). **(V)** Quantification of corrected fluorescent PIP2 levels in cyst cells (CC) depleted of *shi, AP-2α* or *Chc* function. Newly eclosed male flies were shifted at 30°C to activate the RNAi for 4 days. Testes oriented with anterior at left. Image frame (A-H, R-U) 246μ m, (I-P) 123μ m This figure is associated with **Figures S7** and **S8**

Similar to knockdown of Dlg-module components, inhibition of *shi, AP-2α* or *Chc* function resulted in increased levels of dpERK in TJ-positive cyst cell nuclei (Fig. 5M-5P, 5Q) and elevated levels of PIP2 as assessed by the PLCδ-PH-GFP reporter (Fig.5R-5U, 5V). Double knockdown of *dlg* with *shi, AP-2α* or *Chc* led to similar phenotypes as the single knockdowns (Fig.S7A-S7F), confirming that double RNAi does not automatically weaken phenotypes.

Lowering EGFR signaling levels by expressing *UAS-EGFR*^*DN*^ in cyst cells (Fig.6B, 6E, 6H) or by removing one copy of the null *EGFR*^*K*^*05115*^^ allele (Fig. 6C, 6F, 6I), partially rescued the phenotypes caused by depletion of *shi, AP-2α* or *Chc* function in cyst cells (Fig.6K). Rescue experiments were scored according to four phenotypic classes: 1) “dlg LOF” included testes that mimicked the effect of acute 4 and 7 days *dlg* cyst cell knockdowns, with strong decrease in number of germ cells (comparable to Fig.1E, S1A, S1J); 2) “partial rescue/dlg weak” included very weak knockdown phenotypes with only small patches of cyst cell clusters devoid of germ cells (comparable to Fig. S1B-S1D) and germ cells filling up the majority of the testis; 2) “wt” for testes with restored spermatogonial and spermatocyte cysts as in wild type; 3) “EGFR LOF” for all phenotypes that resulted in overproliferation of early spermatogonia resembling strong loss of EGFR signaling. When *EGFR*^*DN*^ was overexpressed in *shi, AP-2α* or *Chc* cyst cell knockdowns, less than 10% of the testes scored showed the CME loss of function phenotype, and many testes showed packed spermatogonia and spermatocytes resembling wild type or overproliferation of spermatogonia resembling EGFR loss of function (Figure 6K; Fig.S9). Removing one copy of the null *EGFR*^*K*^*05115*^^ allele also partially rescued testes in which *shi, AP-2α* or *Chc* function had been knocked down in cyst cells, although the rescue was much weaker than for *EGFR*^*DN*^ overexpression, as expected (Fig. 6K).

**Figure 6:**
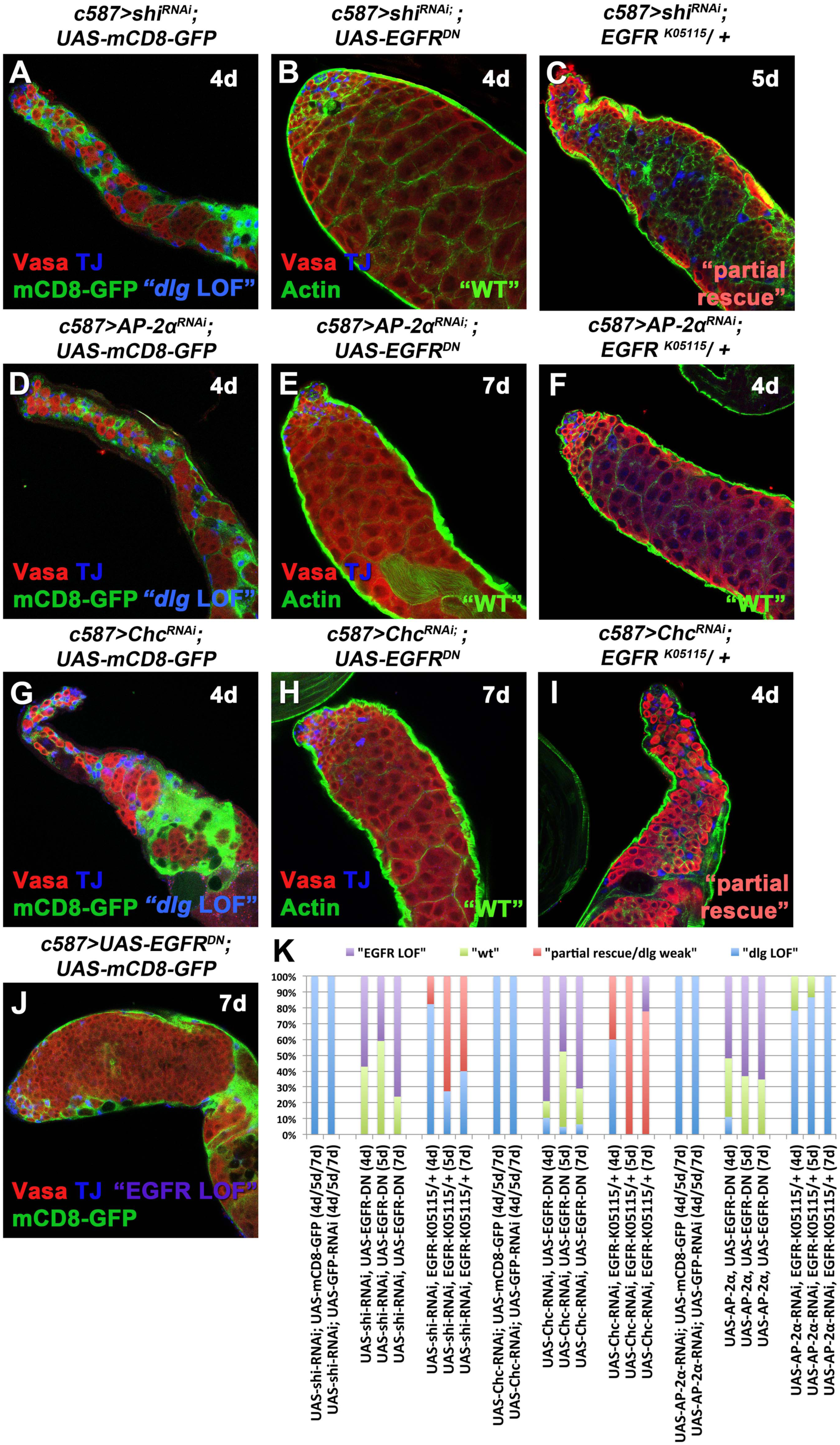
Lowering EGFR signaling levels can rescue clathrin-mediated endocytosis defects in cyst cells. Adult testes of the indicated phenotypes in the background of the *Gal80*^*ts*^. **(A, D, G, J)** *mCD8-GFP* (green) expressed in cyst cells; anti-Vasa (red; germline); anti-TJ (blue; early cyst cells), **(B, C, E, F, H, I)** F-actin stained for phalloidin (green; cyst cells and germline fusome); anti-Vasa (red); anti-TJ (blue). **(K)** Quantifications of the different phenotypic classes accompanying each genotype used, organized in order of phenotypic strength: “dlg LOF”, “dlg weak/partial rescue”, “wt” and “EGFR LOF”. Newly eclosed male flies were shifted at 30°C to activate the RNAi for 4, 5 and 7 days. Testes oriented with anterior at left. Image frame 246μ m. This figure is associated with **Figure S9**

Cyst cell knockdown of Rab11, part of the regulatory machinery for the recycling endosome, led to phenotypes resembling strong EGFR loss of function (Fig. S10E), suggesting that recycling of internalized EGFR back to the plasma membrane may help to maintain proper levels of EGFR activity in early cyst cells. Consistently, knocking down Rab11 in *dlg* depleted cyst cells partially rescued the *dlg* loss of function phenotype (Fig. S10C, S10F), as seen for reducing EGFR signal transduction pathway by other means.

## DISCUSSION

Male gamete development requires close communication between germline and neighboring somatic cells. Such interactions are essential for stem cell maintenance and the coordinated differentiation of both the germ cells and their somatic support cells. Our results indicate that both the cortical cell polarity and scaffolding proteins Dlg, Scrib and Lgl, and components of clathrin-mediated endocytosis (CME) play an important role in the intimate germline-cyst cell signaling exchange that sets up a functional microenvironment required for proper differentiation of early germ cells. Our findings show that loss of *dlg, scrib, lgl* or CME components in the cyst cell lineage results in phenotypes resembling overactivation of the EGFR signal transduction pathway, including non-cell-autonomous effects on germ cells leading to death of late spermatogonia or early spermatocytes. Lowering activity of the EGFR signaling components in cyst cells partially rescued and restored those defects, indicating that the Dlg-module and CME can fine tune EGFR pathway activity in cyst cells, either through effects on signal transduction components or internalization of the receptor (Fig. 7).

**Figure 7:**
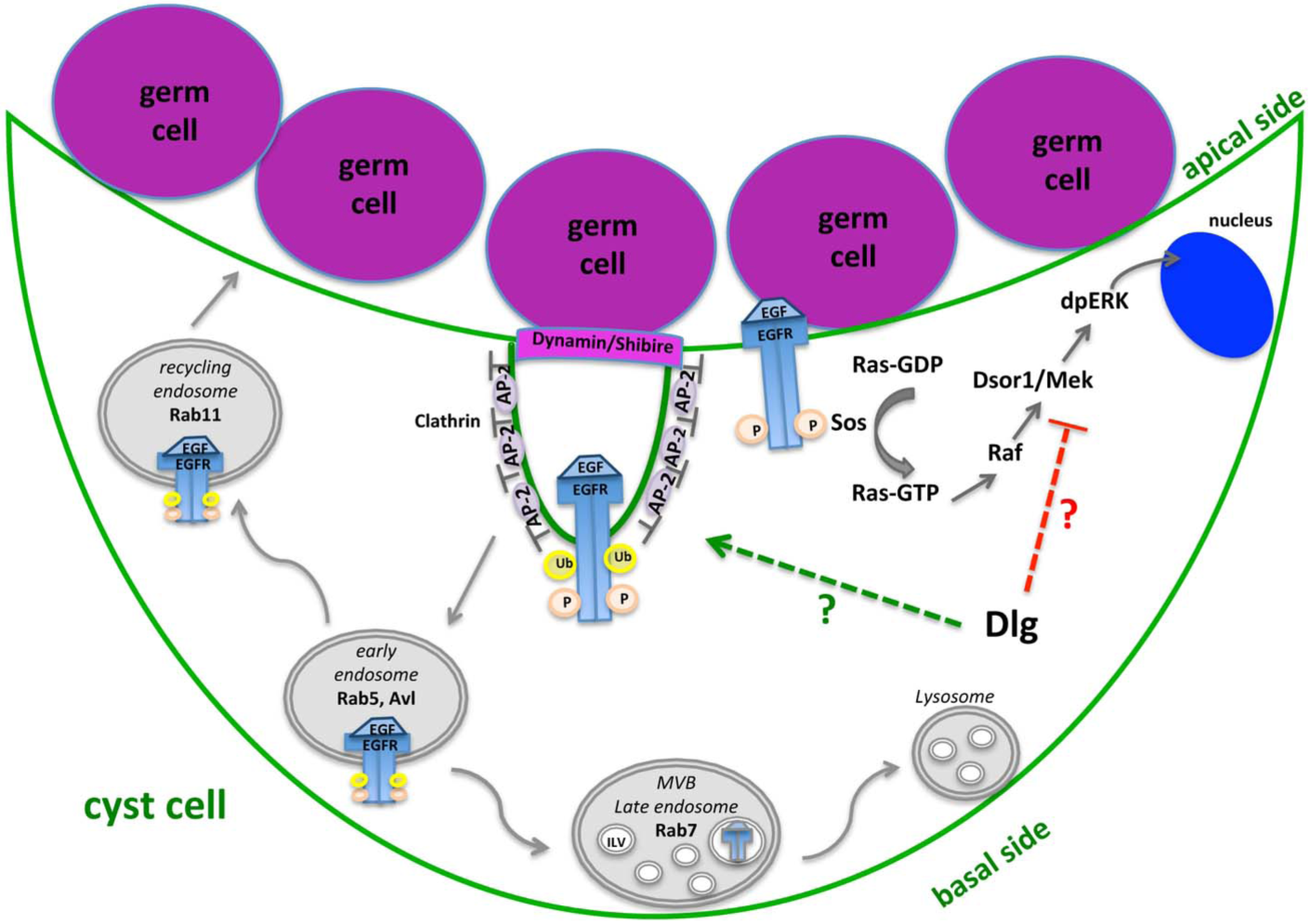
Model diagram showing the EGFR signaling pathway and clathrin-mediated EGFR endocytosis in cyst cells, and the possible involvement of the Dlg-module. Activation of EGFR at cyst cell membranes upon binding of the EGF ligand Spitz secreted from the germ cells activates the Ras/MAPK cascade that leads to dpERK entering the cyst cell nucleus. Clathrin-mediated endocytosis (CME) attenuates EGFR signaling by endocytosis of activated EGFR from the membranes. Once in early endosomes, EGFR can either recycle back to the membrane via Rab-11 containing recycling endosomes or get into multivesicular endosomes (MVE) to be targeted for degradation. Dlg-module components could downregulate EGFR signaling either by cooperating with CME components or by binding and inactivating Ras/MAPK signaling components.

Sequential steps in early male germ cell differentiation in *Drosophila* have been shown to require increasing levels of EGFR activation in somatic cyst cells (Hudson et al., 2013). Receipt of EGF signaling from the germline promotes encapsulation of the gonialblast by two cyst cells (Sarkar et al., 2007; Schulz et al., 2002) and consequent activation of the EGFR on cyst cells is required for germ cells to properly enter the transit-amplifying program of synchronous mitotic divisions (Kiger et al., 2000; Tran et al., 2000). Higher levels of EGFR activation in cyst cells are required for spermatogonia to end the TA divisions and initiate the spermatocyte program (Hudson et al., 2013). It is possible that the death of late spermatogonia/early spermatocytes we observe may be due to cyst cells experiencing very high sustained levels of EGFR activation sending mixed or stage inappropriate signals to the germ cells they enclose.

Attenuation of EGFR signaling via endocytosis has been demonstrated in many tissues and organs across species, and deregulation of this process has been implicated in cancer initiation and progression (Conte and Sigismund, 2016; Dobrowolski and De Robertis, 2011). Upon binding of EGF ligand, the EGFR becomes activated by phosphorylation and recruits adaptor proteins that stimulate the Ras/MAPK cascade. However activation by ligand binding also stimulates internalization of the EGFR receptor, mainly by clathrin-mediated endocytosis (Conte and Sigismund, 2016; Czech, 2000; Dobrowolski and De Robertis, 2011; Goh and Sorkin, 2013; Krahn and Wodarz, 2012). Activated EGFR is recruited to clathrin-coated pits by interacting with the adaptor protein (AP)-2 complex, then clathrin-coated pits invaginate and pinch off with the action of the GTPase Dynamin, encoded by the *shibire* gene in *Drosophila* (Chen et al., 2002; Conte and Sigismund, 2016; Dobrowolski and De Robertis, 2011; Goh and Sorkin, 2013). Knockdown of any of these CME components in the cyst cell lineage resulted in phenotypes resembling those caused by forced overactivation of the EGFR or Ras in cyst cells, suggesting that CME is required to maintain levels of EGFR signaling in a range appropriate for the normal developmental progression of male germline cysts. Much of the activated EGFR internalized by endocytosis is targeted for degradation, while depending on the cell type, some is recycled back to the plasma membrane. Our data that loss of function of Rab11 in cyst cells can rescue *dlg* loss of function defects raises the possibility that recycling of the EGFR back to the surface dependent on Rab 11 plays a role in keeping EGFR signaling activity in a physiological range appropriate for the germ cell cyst stage.

The Dlg module could act to attenuate signaling via the EGFR in cyst cells either by potentiating endocystosis of the receptor or, independently of CME, by down regulating activity of signal transduction mechanisms downstream of the EGFR. Numerous studies support cooperation of the Dlg-module with vesicle and membrane trafficking, including endocytosis, exocytosis, recycling of endosomes to the cell membrane and retrogade trafficking, (shuttling between endosomes, biosynthetic, or secretory compartments) (Elsum et al., 2012; Humbert, 2015; Stephens et al., 2018). In *Drosophila*, Lgl controls endocytosis of the Notch regulator Sanpodo (Roegiers et al., 2005) in sensory precursor cells and attenuates Notch signaling. Also, in eye disc epithelia, Lgl associates with early to late endosomes and lysosomes and attenuates Notch signaling by limiting vesicle acidification (Parsons et al., 2014), which is achieved by promoting binding of vATPase to Vap33 (Portela et al., 2015). Likewise, Scrib optimizes BMP signaling by regulating the basolateral localization of the BMP receptor Thickveins and its internalization in Rab5-positive endosomes in wing epithelia (Gui et al., 2016). In mammalian MDCK (Madin-Darby canine kidney) epithelial cells, Scrib negatively regulates retromer-mediated E-cadherin (E-cad) trafficking to the Golgi (Lohia et al., 2012). Several mechanisms also link the Dlg-module to regulation of signal transduction along the EGFR/MAPK pathway [reviewed in detail in (Humbert, 2015; Milgrom-Hoffman and Humbert, 2017; Stephens et al., 2018)]. For example, human Scrib (hScrib) binds ERK (MAPK) (via its PDZ1 domain) and anchors it to membrane sites to prevent ERK phosphorylation, thus inhibiting signaling via Ras (Nagasaka et al., 2010). Scrib also influences EGFR pathway signaling by binding the Arf-GAP/βPix (Pak-interactive exchange factor) that acts as a MEK-ERK scaffold (Audebert et al., 2004). Moreover, mammalian Dlg1 binds phosphorylated MEK2, which phosphorylates and activates ERK, and regulates its cortical and perinuclear localization (Gaudet et al., 2011; Maiga et al., 2011), while Dlg2, Dlg3 and Scrib interact with PP1 phosphatases to downregulate ERK phosphorylation (Nagasaka et al., 2013). Presumably, the Dlg-module regulates signaling by cooperating with distinct cellular trafficking components in different tissues in part because the availability of partner proteins varies significantly based on cell type and developmental stage.

We also observed that levels of the membrane phospholipid PIP2 appeared elevated in cyst cells in which function of Dlg module components had been knocked down or constitutively active EGFR or Ras had been forcibly expressed, and that loss of function of *Skittles*, the PIP5 Kinase that makes PIP2, partially alleviated the effects of loss of function of Dlg-module components, suggesting that the levels of PIP2 may contribute to the phenotype. Membrane PIP2 is a critical regulator of both membrane trafficking and cell signaling components, including the EGFR (Marat and Haucke, 2016). PIP2 binds to the EGFR juxtamebrane domain (Abd Halim et al., 2015) and enhances EGFR phosphorylation and activation (Michailidis et al., 2011). Down-regulation of PIP2 levels via pharmacological inhibition or manipulation of PIP5K levels reduced EGFR tyrosine phosphorylation (Michailidis et al., 2011), consistent with our finding of partial rescue by knocking down function of *Skittles*/PIP5K. It is also possible that EGFR activation promotes the synthesis of PIP2 by the PIP5K kinase, as enhanced EGFR activity in response to EGF resulted in increased synthesis of PIP2 mediated by Arf6 in bovine brain cytosol (Honda et al., 1999) and increasing evidence indicates that PIP5K functions as a downstream effector of Arf6 to regulate cellular functions such as exocytosis, endocytosis, endosomal recycling, and membrane ruffle formation (Funakoshi et al., 2011). Following vesicle endocytosis by CME, maturation of clathrin-coated pits initiates a cascade that locally converts PIP2 to different phosphatidylinositols and finally to PI(3)P, an essential feature of early endosomes, in a cascade that is important for vesicle uncoating (Marat and Haucke, 2016; Posor et al., 2015). Blocking early steps of CME by knocking down function of Shibire AP-2α, or Clathrin heavy chain may thus allow accumulation of higher levels of PIP2 in cyst cell plasma membranes, where it can promote EGFR activation.

An alternative possibility is that Dlg, Scrib and Lgl regulate polarity of the squamous epithelial-like cyst cells, much as they regulate apical/basal polarity in columnar epithelia. If so, and if polarized distribution of the EGFR is important for proper levels of signal reception in cyst cells, as in many polarized epithelial cells (Kuwada et al., 1998; Singh and Coffey, 2014; Vermeer et al., 2003), loss of cyst cell polarity with knockdown of Dlg module components may affect access of the EGFR to ligand Spitz secreted by the germ cells and thereby attenuate the EGFR signaling.

Our results provide new insights into the role of Dlg, Scrib and Lgl in regulating signaling pathways and uncovers new regulatory strategies on how endocytosis and signaling cooperate with polarity cues to coordinate stem cell maintenance vs. differentiation and sculpt developing tissues. The high degree of conservation of Dlg, Scrib, Lgl across species allows us to drive qualitative conclusions on how stem cells communicate with their neighbors to replenish differentiated tissues and to extend the basic mechanistic features on cell-cell communication uncovered here, to other tissues and organisms.

## EXPERIMENTAL PROCEDURES

### Fly stocks and husbandry

Fly stocks used in this study are described in FlyBase (www.flybase.org). A detailed description of fly stocks used in this study is provided in “Supplemental Experimental Procedures”. All *UAS-gene*^*RNAi*^ stocks are referred to in the text as *gene*^*RNAi*^ for simplicity reasons. Knockdowns were performed using the *UAS-GAL4* system by combining the *UAS-RNAi* fly lines with the cell-type specific *c587-GAL4* driver and *αtub84B-Gal80*^*ts*^. Crosses were raised at 18°C until adult flies hatched. Then males with the correct genotype were shifted at 30°C for 2-7 days depending on the experimental needs and the phenotypes were analyzed.

### Immunofluorescence staining and microscopy

Whole mount testes immunostaining was performed as previously described (Papagiannouli and Mechler, 2009). For testes immunostaining in the presence of GFP, 1% PBT (1% Tween-20 in PBS) was used instead of 1% PBX in all steps. For the TUNEL assay, tissue was processed with the standard protocol except that after fixation, the protocol from In Situ Cell Death Detection Kit (TMR Red, Sigma/Roche) was followed. For a detailed staining protocol and a detailed list of antibodies used in this study see “Supplemental Experimental Procedures”.

Confocal images were obtained using a Leica SP8 system (CSIF Beckman Center, Stanford University School of Medicine; CECAD, University of Cologne). Pictures were finally processed with Adobe Photoshop 7.0. Quantifications were done using FiJi/ImageJ by measuring “Corrected Total Cell Fluorescence” (CTCF) [CTCF = Integrated Density - (Area of selected cell X Mean fluorescence of background readings)].

## AUTHOR CONTRIBUTIONS

F.P. designed, performed, interpreted experiments, wrote the paper and obtained funding to support the study. C.W.B. did preliminary work on EGFR. M.T.F. co-designed and interpreted experiments, assisted with writing the paper and obtained funding to support the study.

## ACKNOWLEDGMENTS

We thank Jaclyn Lim and Susanna Brantley for preliminary data, fly stocks, experiment interpretation and helpful discussions. We would like to thank the *Drosophila* community for providing us generously with fly stocks and antibodies, in particular Dorothea Godt, Ruth Lehmann, Antoine Guichet, Dennis McKearin, the Developmental Studies Hybridoma Bank, Vienna *Drosophila* Resource Center (VDRC) and Bloomington *Drosophila* Stock Center. We would like to deeply thank Maria Leptin, as part of this work was done in her lab. We apologize to all whose work was not sited due to space limitations. This work was supported by the DFG/ PA2659/3-1 to F.Papagiannouli and NIH grant 1 R01 GM080501 to M.T. Fuller. We thank the Cell Sciences Imaging Facility (Beckmann Center, Stanford University) and the CECAD Imaging Facility (University of Cologne) for their technical support.

## Abbreviations

CySCs: somatic cyst stem cells
CCs: cyst cells
GSCs: germline stem cells
*dlg*: *discs large*
*scrib*: *scribble*
*lgl*: *lethal (2) giant larvae*
CME: clathrin-mediated endocytosis
*Chc*: *clathrin heavy chain; shi, shibire*
PIP2: PtdIns(4,5)P2.

